# Insights into H_2_O_2_-induced signaling in Jurkat cells from analysis of gene expression

**DOI:** 10.1101/2022.10.20.513094

**Authors:** Megan F. Taylor, Michael A. Black, Mark B. Hampton, Elizabeth C. Ledgerwood

## Abstract

Hydrogen peroxide (H_2_O_2_) is a ubiquitous oxidant produced in a regulated manner by various enzymes in mammalian cells. H_2_O_2_ reversibly oxidises thiol groups of cysteine residues to mediate intracellular signalling. Whilst examples of H_2_O_2_ dependent signalling have been reported, the exact molecular mechanism(s) of signalling and the pathways affected are not well understood. Here, the transcriptomic response of Jurkat T cells to H_2_O_2_ was investigated to determine global effects on gene expression. With a low H_2_O_2_ concentration (10 μM) that did not induced an oxidative stress response or cell death, extensive changes in gene expression occurred after 4 hours (6803 differentially expressed genes). Of the genes with greater then 2-fold change in expression, 85% were upregulated suggesting that in a physiological setting H_2_O_2_ predominantly activates gene expression. Pathway analysis identified gene expression signatures associated with FOXO and NTRK signalling. These signatures were associated with an overlapping set of transcriptional regulators. Overall, our results provide a snapshot of gene expression changes in response to H_2_O_2_, which, along with further studies, will lead to new insights into the specific pathways that are activated in response to endogenous production of H_2_O_2_, and the molecular mechanisms of H_2_O_2_ signalling.

## 1. Introduction

Hydrogen peroxide (H_2_O_2_) can mediate signal transduction by reversible oxidation of redox-sensitive cysteine residues in target proteins, and is considered a major reactive oxygen species for redox regulation (2–7). H_2_O_2_ is a suitable second messenger as it is produced in a regulated manner, is small and can diffuse through cell membranes, and is specific in its reactions with thiols and transition metal centres (4,5). While over 41 enzymes can produce H_2_O_2_ or its precursor O_2^.-^_, the two main sources of H_2_O_2_ are membrane bound NADPH oxidase (NOX) enzymes and the mitochondrial electron transport chain (6). NOX enzymes are activated in response to a variety of signals including growth factors and cytokines. They transfer electrons from intracellular NAD(P)H across plasma or endosomal membranes to molecular oxygen generating superoxide which is then converted to H_2_O_2_ (8,9). The H_2_O_2_ enters the cytosol via passive diffusion or facilitated transport by aquaporins to induce a signalling response via direct or indirect protein oxidation. Mitochondrial ROS production is predominantly via reverse electron transport at complex I, and from reaction of oxygen with ubisemiquinone at complex III, generating superoxide (and subsequently H_2_O_2_) in either the mitochondrial intermembrane space or the mitochondrial matrix (10).

H_2_O_2_ is a pleiotropic molecule that has been reported to affect multiple signalling pathways. For example, it is well documented that in cells stimulated with growth factors, such as epidermal growth factor (EGF), production of H_2_O_2_ results in oxidation of a catalytic cysteine residue of protein tyrosine phosphatases (PTPs) leading to their inactivation (11,12). Inactivation of PTPs is required concurrent with the activation of protein tyrosine kinases (PTKs) for the propagation of an intracellular signal. Often oxidation of specific targets leads to alterations in phosphorylation cascades and subsequent activation of downstream transcription factors leading to changes in gene expression. For example, the MAPKKK ASK1 that activates either p38 or JNK is redox regulated (13–15). In response to H_2_O_2_ treatment, ASK1 forms multimeric disulphide linked species which are essential for full JNK activation, nuclear translocation and changes in transcription (14).

There is an abundance of literature implicating H_2_O_2_ in cellular signalling, predominantly focussing on investigating specific signalling targets. In addition, microarrays and RNA-Seq have been used to provide a global overview of the transcriptomics response of budding and fission yeasts, and cultured human cells, to H_2_O_2_ (16–29). Collectively these studies support a role for H_2_O_2_ in controlling the expression of a range of genes, including p53 target genes, apoptosis genes and signal transduction genes. However, in many of these studies it is unclear whether changes in gene expression are due to the direct impact of H_2_O_2_, because the concentrations used induce cell death and/or the timepoint studied is many days after addition of H_2_O_2_ to the cells.

Here, we have used a discovery approach to investigate gene expression changes in response to physiologically relevant concentrations of H_2_O_2_. The physiological concentration of H_2_O_2_ is estimated to lie between 1-100 nM and to rise transiently to 500-700 nM under oxidative signalling events (30,31). We performed RNA-Seq analysis on RNA extracted from human Jurkat T cells four hours after treatment with 10 μM H_2_O_2_ and observed widespread changes in gene expression. Using Reactome Pathway Analysis, we identified several clusters of genes involved in intracellular signalling suggesting that H_2_O_2_ may play a role in activation of these pathways. Further analysis of these molecular signalling events may provide insight into the role of endogenously produced H_2_O_2_ in cell signalling.

## 2. Materials and Methods

### 2.1 Cell culture and H_2_O_2_ treatment

Jurkat T lymphoma cells (TIB-152 clone E6-1 from ATCC) were maintained at a density of less than 1 x 10^6^ cells/mL in RPMI 1640 (Gibco, Invitrogen) with sodium bicarbonate, 10% foetal bovine serum (FBS), and penicillin/streptomycin supplements (Sigma). Cells were maintained at 37 °C in a humidified atmosphere containing 5 % CO_2_. Cell line identity was confirmed by STR analysis and cultures were regularly screened for mycoplasma contamination.

H_2_O_2_ concentration was determined spectrophotometrically (*ϵ*_240_ = 43.6 M^-1^ s ^-1^). For treatment with H_2_O_2_, cells were harvested by centrifugation and resuspended in PBS at a density of 1 x 10^6^ cells/mL to prevent H_2_O_2_ interacting with components in RPMI media and to slow the reduction of oxidised proteins (32). Following treatment, cells were pelleted, washed in 1 mL PBS, resuspended in 1 mL TRIzol (Life Technologies), and stored at −80 °C.

### 2.2 Analysis of cell death

Jurkat cells were resuspended at a density of 1 x 10^6^ cells/mL in 2 mL PBS and equilibrated to 37 °C/5 % CO_2_ before treatment with 0, 10, 20 or 500 μM H_2_O_2_ per 10^6^ cells. After incubation at 37 °C for 4 hours, the cells were pelleted, and resuspended in 2 mL RPMI containing 10 % FBS. After 24 hours at 37 °C/5 % CO_2_, the cells were pelleted then washed twice in 1 mL PBS. Subsequently, the cells were resuspended in 100 μL PBS with 4 μL of propidium iodide (PI) stain (50 μg/mL). The cells were incubated in the dark for 10 minutes before analysis by flow cytometry using the Guava^®^ easyCyte^™^ 5HT HPL Benchtop flow cytometer (Merck). Data were analysed using FlowJo^™^ v10.2 Software (BD Life Sciences).

### 2.3 RNA sequencing

RNA extractions were carried out using TRIzol according to the manufacturer’s instructions. RNA quality was assessed by running an RNA 6000 Nano chip on a 2100 Bioanalyser (Agilent). TruSeq (Illumina) libraries for polyA+ RNA-Seq were prepared from 250 ng RNA per sample using the TruSeq RNA Library Preparation Kit v2. The libraries were multiplexed (24 samples per pool) and sequenced to obtain 50 bp single-end reads on the Illumina HiSeq 3000 platform (UCLA Technology Center for Genomics & Bioinformatics). Library preparation and sequencing was performed in two batches, each with four biological replicates of control and treated cells.

### 2.4 Read mapping and differential expression analysis

The quality of the raw data was evaluated using FastQC (33) and reads were mapped to human genome version GRCh38 using STAR v2.7.1a (34), resulting in an overall mapping efficiency of > 80 %. The GENCODE hv35 was used as the annotation file and the reads per gene were counted using the HTseq package v0.6.1 (35), with parameters -m union (recommended) -s no. Differential gene expression was performed with limma (36,37). Genes with low expression in the samples (< 160 reads across 16 samples) were excluded prior to differential expression analysis. Principal Component Analysis (PCA) and the heatmap.2 function in the gplots package (38) for R (v4.0.2) were used to visualise similarities and differences among the control and treated samples (39). The PCA plot of the samples showed that approximately 70% of the variation in the data was explained by H_2_O_2_ treatment, however, the remaining variation was a batch effect associated with the two separate library preparations and sequencing runs (Figure S1). For this reason, batch correction was completed using limma. Adjusted p-values were calculated using the Benjamini–Hochberg method (40) and a value of 0.05 was set as threshold for statistical significance of differential gene expression. A volcano plot to visualise differential gene expression was generated using the R package EnrichedVolcano (41).

### 2.5 Gene Ontology and pathway analysis

Significantly differentially expressed gene ensemble IDs were converted to entrez IDs using the AnnotationDb package (42) in R. Gene set analysis was performed using GOseq (43) with Reactome pathways (44) to correct for gene length, as well as via a non-length corrected hypergeometric model. The overlap between the output from GOSeq and the hypergeometric model suggested there was little or no length bias in the data. Therefore, gene set enrichment analysis (GSEA) (45) was performed for Reactome pathways (44) using the ReactomePA (v1.9.4) (46) package in R. Of the 6803 DEGs, 3356 were suitable for pathway analysis (2780 were not annotated with a Reactome pathway ID and the remainder didn’t have a corresponding Entrez ID). The heatmap.2 function was used to plot the proportion of overlapping gene content between pathways that were found to be significantly enriched in the data.

### 2.6 RT-qPCR

Selected differentially expressed genes were analysed by RT-qPCR. RNA (1 μg) was reverse transcribed to cDNA using SuperScript IV VILO Master Mix with ezDNAse enzyme (ThermoFisher) according to the manufacturer’s instructions. One-fifth of the volume of cDNA reaction mixture was used in the qPCR reaction containing 1 x SYBR Green Master Mix (ThermoFisher) in a 384-well-plate. Cycling conditions were as follows: 2 min at 50 °C and 2 min at 95 °C then 40 cycles of amplification at 95 °C for 15 s and 60 °C for 1 min. Primer specificity was initially tested using agarose gel electrophoresis and was subsequently monitored using melting curve analysis. RPL27 (the most stable of five candidate reference genes analysed using Normfinder (47)) was used as the reference gene and gene expression relative to RPL27 was calculated as 2^-ΔCt^ (48). Primer information is provided in Table S1.

### 2.7 Motif Analysis using ChEA3 and TRAP

Two algorithms, TRAP (Transcription Factor Affinity Prediction) and ChEA3 (ChIP-X Enrichment Analysis 3) were used to identify transcription factors which may be responsible for regulating gene sets identified by pathway analysis. TRAP uses a biophysical model to predict which transcription factors have the highest affinity for a given set of DNA sequences (49,50). TRAP (multiple sequences) at http://trap.molgen.mpg.de/cgi-bin/home.cgi was used. The 1000 bp upstream flanking region of each significantly differentially expressed gene in each gene set underlying identified Reactome Pathways was generated using BioMart and saved as a .fasta file. This file was submitted to TRAP and enriched binding motifs were predicted and compared with random promoters using the settings ‘jaspar_vertebrates’ and ‘human_promoters’. Motifs for significant hits were obtained from JASPAR CORE

Vertebrates clustering (https://jaspar.genereg.net/matrix-clusters/vertebrates/?detail=true). ChEA3 ranks TFs based on the overlap between gene input and annotated sets of TF targets in background databases (available at https://maayanlab.cloud/chea3/) (51). Gene symbols for each each gene set to be analysed were submitted. As a control, the sample function in R was used to generate random set of 1050 human genes from the genes expressed in Jurkat cells. Transcriptional regulators identified that had low expression in the Jurkat cells (< 160 reads across 16 samples) were removed from the results.

### 2.8 Statistical analysis

For analysis of cell death, changes in variables were analysed by one-way ANOVA with Dunnett’s multiple comparison test assuming single pooled variance (GraphPad Prism v9.3.1). For analysis of qPCR data, changes in variables were analysed by two-way ANOVA with Tukey’s post-hoc test for multiple comparisons (GraphPad Prism v9.3.1). In both cases, differences were considered statistically significant at *P* < 0.05.

## 3. Results

The majority of studies investigating the transcriptomic response to H_2_O_2_ in mammalian cells have focused on conditions that induce a stress response and/or monitor the impact after several days. Here we investigate the early gene expression response to a bolus treatment with H_2_O_2_ at a concentration which modelling suggests will result in low nM intracellular H_2_O_2_ (31).

### 3.1 Treatment with 10 μM H_2_O_2_ does not induce cell death

In Jurkat cells, H_2_O_2_ is reported to induce apoptotic or necroptotic cell death at concentrations of 25 - 200 μM and greater than 200 μM respectively (52). At 10 μM H_2_O_2_/10^6^ cells/mL there was no significant cell death at 24 h, and only a small increase with 20 μM H_2_O_2_/10^6^ cells/mL (Figure 1). As expected, treatment with 500 μM H_2_O_2_/10^6^ cells/mL killed almost all the cells. Therefore treatment with 10 μM H_2_O_2_/10^6^ cells/mL and harvesting after 4 h was used for analysis of gene expression (referred to as 10 μM hereafter).

**Figure 1:**
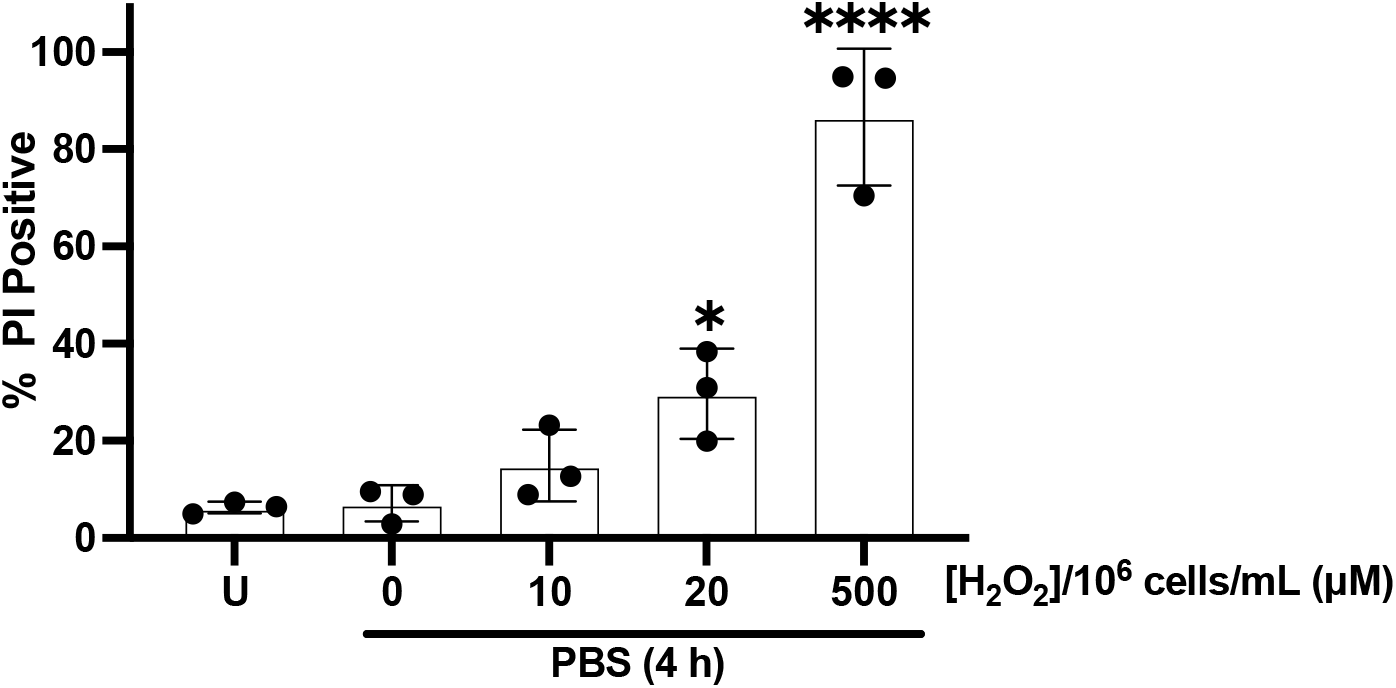
Cell death of Jurkat T cells after treatment with varying concentrations of H_2_O_2_. Cells were treated with H_2_O_2_ in PBS and after four hours returned to RPMI/FCS for a further 24 h before staining with PI and analysis by flow cytometry. U = cells that remained in RPMI/FCS for the full 28 h. \**P* <0.05, \*\*\*\**P* < 0.0001 compared to U (n=3 ± SD).

### 3.2 H_2_O_2_ treatment of Jurkat cells leads to changes in gene expression

RNA-Seq was performed on RNA isolated from control cells, and cells harvested 4 h after treatment with 10 μM H_2_O_2_, with 8 biological replicates for each. Of the ~15,000 genes identified by RNA Seq, 6,803 were significantly differentially expressed genes (DEG, defined by *P*_adj_ < 0.05), the majority with low fold-change (Figure 2). When considering the 1,050 DEG with greater than 2-fold change in expression (| log_2_ fold-change | (| log_2_FC |) >1), 86% had increased expression. There was limited induction of oxidative stress response genes (as defined by Desaint et al (20)) in our dataset (all | log_2_FC | < 0.3). Overall, this indicates that at a low concentration H_2_O_2_ predominantly induces gene expression.

**Figure 2:**
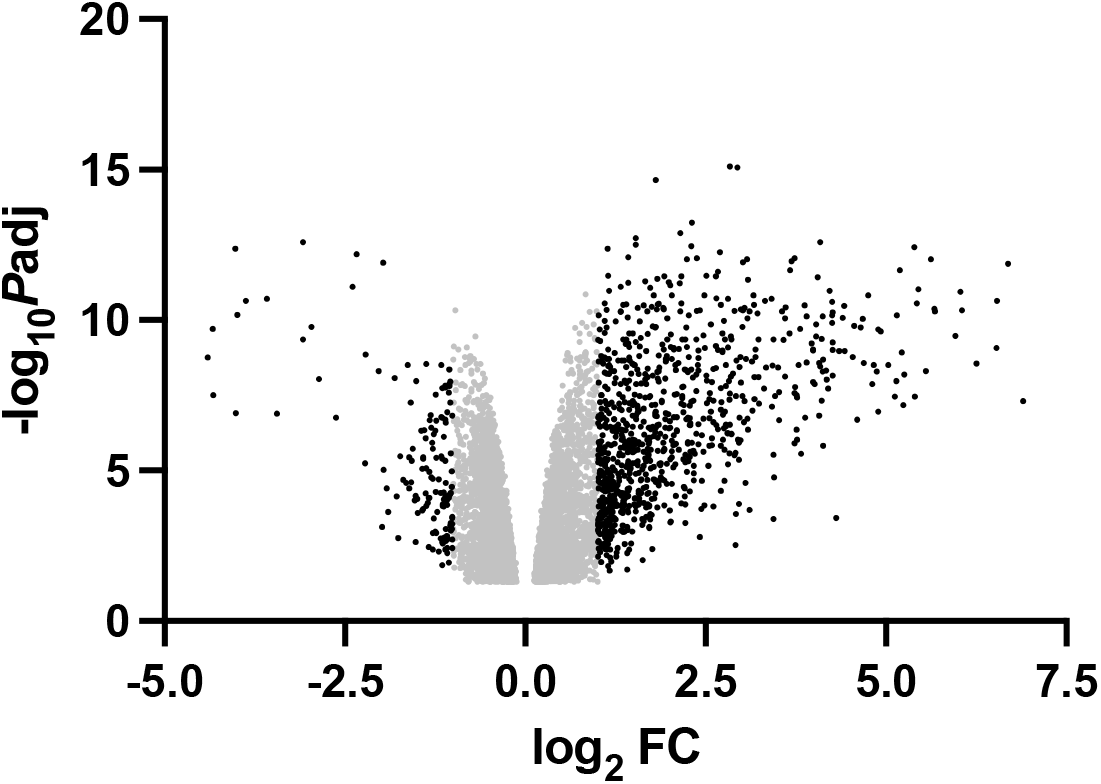
Volcano plot of differentially expressed genes following H_2_O_2_ treatment. The black dots are DEG with |log_2_FC| > 1 and the grey dots are DEG with −1 ≤ log_2_FC ≤ 1. The list of DEG is in Table S2.

### 3.3 Reactome Pathway Analysis identified gene sets which respond to H_2_O_2_

To identify pathways enriched in the list of DEG, and thereby provide insight into the biological impact of H_2_O_2_-mediated signalling, we used the *Homo sapiens*-focused Reactome Knowledgebase (44). Focusing on the 1050 genes with >2-fold change in expression, 391 were unique and able to be mapped in Reactome (Table S2). This analysis identified 21 Reactome pathways with a range of significance (*P*_adj_ from 8.3×10^-6^ to 0.04) (Figure 3A). As expected from the analysis of cell death and expression of oxidative stress genes, none of the identified pathways were within the superpathways “Programmed Cell Death” or “DNA Repair”, confirming that 10 μM H_2_O_2_ induces a signalling response independent of a classic oxidative stress response. An additional 330 Reactome pathways were identified using the mappable genes from the full set of DEG (Table S4).

**Figure 3:**
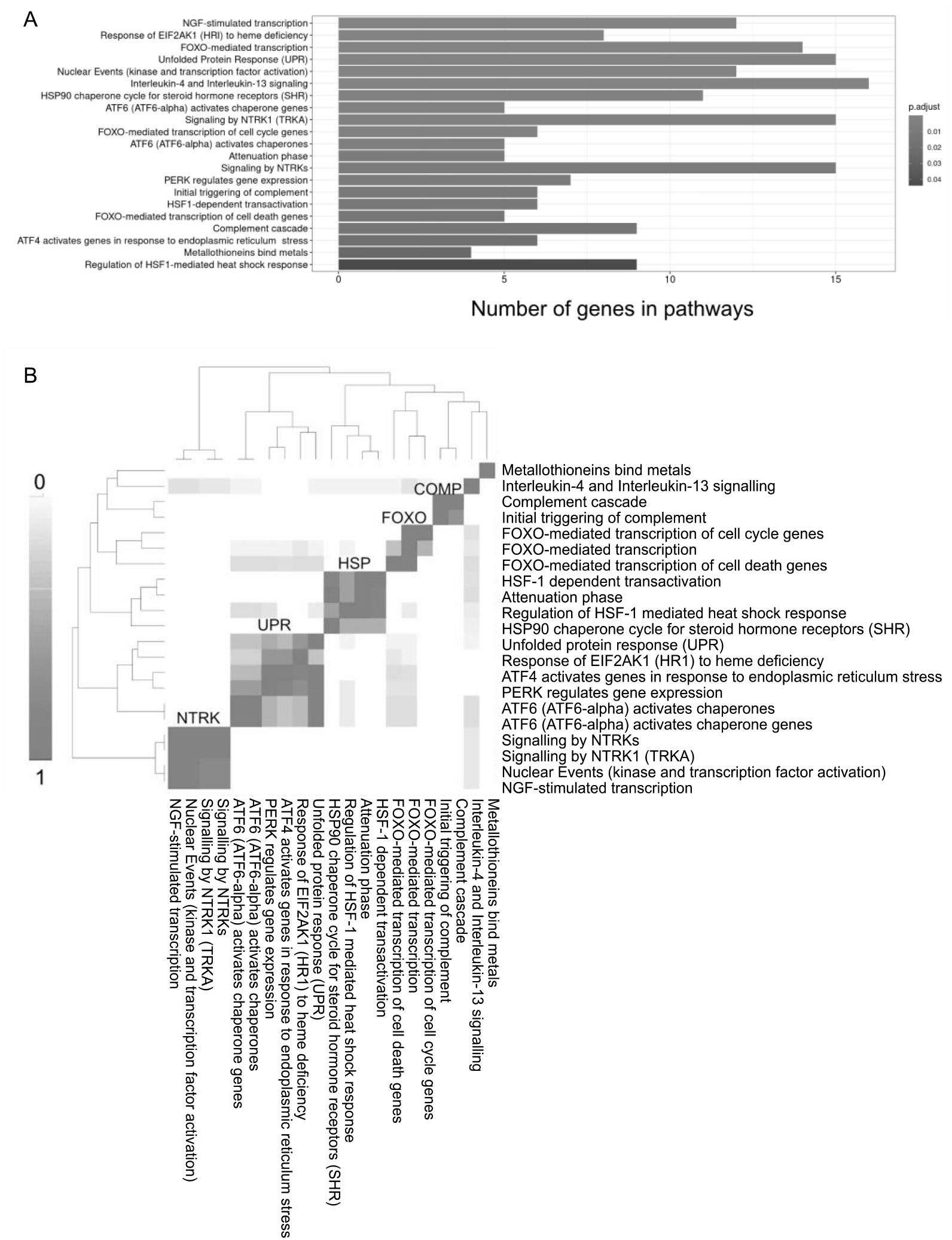
Enriched Reactome pathways in Jurkat T cells treated with H_2_O_2_. Pathway analysis was completed using genes with *P*_ad_j < 0.05 and > 2-fold change in expression. **A.** Bar graph is ordered based on *P*adj values (highest to lowest), and sizes of bars indicates the number of DEGs within each pathway. Table S3 contains more detailed information for each pathway. **B.** Plot showing the overlap in gene content between each Reactome pathway regulated by H_2_O_2_. The five clusters are annotated as NTRK, UPR, HSP, FOXO and COMP. The values in the plot are computed by the number of genes appearing in both pathways (A and B) as a proportion of the total size of either pathway A (upper diagonal) or pathway B (lower diagonal). The scale is from 1 (100% gene overlap between Reactome pathways) to 0 (0% gene overlap between Reactome pathways).

There is frequently gene membership overlap between Reactome pathways (53). Therefore a plot showing the overlap in gene content between pathways was used to gain a more precise understanding of biological consequences of H_2_O_2_ signalling (Figure 3B). Of the 21 significant Reactome pathways, 17 grouped in five clusters: NTRK signalling, FOXO signalling, unfolded protein response (UPR), heat shock response, and complement (Comp). Two Reactome pathways, “metallothioneins bind metals” and “IL4 & IL14 signalling”, exhibited only minimal gene-sharing with other enriched pathways.

### 3.4 FOXO signalling and NTRK signalling are H_2_O_2_-dependent

Reactome pathways that respond to H_2_O_2_ were identified, however, there was a caveat to this result. The H_2_O_2_-treated RNA samples were harvested from Jurkat cells incubated in PBS for four hours. Therefore, the pathways identified may be H_2_O_2_-dependent and/or PBS-dependent. To determine which responses were H_2_O_2_ dependent, and to gain insights into the pattern of activation, RT-qPCR of two genes from each of the NTRK, FOXO and UPR clusters was carried out over a time-course at 0, 5, 10 and 20 μM H_2_O_2_.

Expression of the NTRK cluster genes *FOS* and *FOSB* was H_2_O_2_ dependent, with expression increasing over time in response to 10 and 20 μM H_2_O_2_ (Figure 4A). The FOXO cluster genes *BTG1* and *CITED2* showed similar H_2_O_2_-dependent increases over time at 10 and 20 μM H_2_O_2_ (Figure 4B). There was no change in expression of these four genes in the absence of H_2_O_2_. In contrast the expression of the UPR cluster genes *BIP* and *CHOP* was not dependent on H_2_O_2_, confirming that the UPR is activated due to culture in PBS (Figure 4C). Interestingly, 20 μM H_2_O_2_ appeared to protect the cells from PBS-induced UPR activation. Overall these results confirm that H_2_O_2_ induces dose- and time-dependent changes in gene expression consistent with activation of FOXO and NTRK signalling.

**Figure 4:**
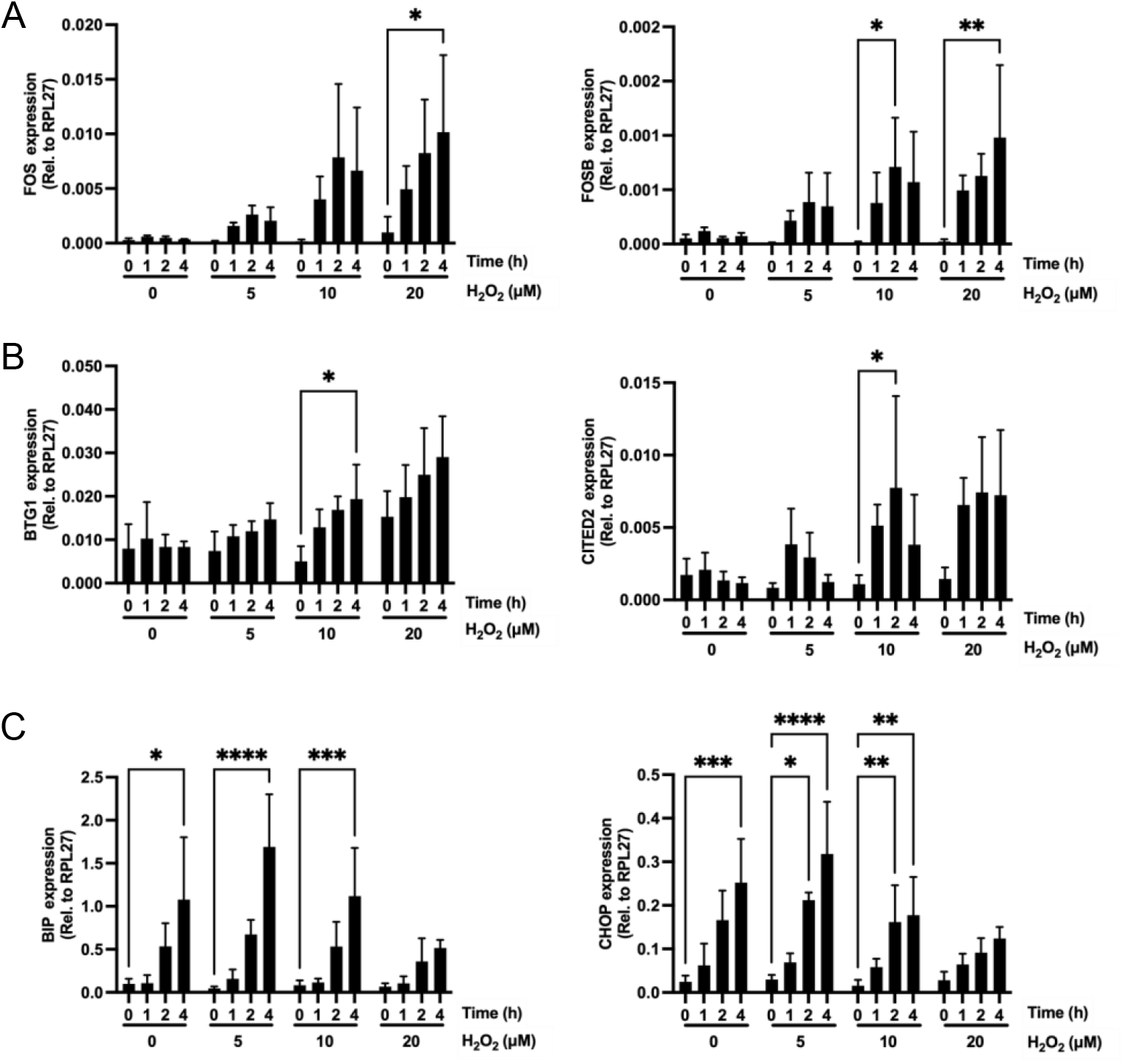
RT-qPCR showing changes in expression of selected genes from Reactome Pathways. **A.** NTRK cluster genes *FOS* and *FOSB*. **B.** FOXO cluster genes *BTG1* and *CITED2*. **C.** UPR cluster genes *BIP* and *CHOP*. For each concentration of H_2_O_2_ a two-way ANOVA with multiple comparisons testing was used to determine statistical significance. Note that 0 h for each gene is the same data. \**P* <0.05, \*\**P* < 0.01 and \*\*\*\**P* < 0.0001 (n = 3-12±SD). The results at 4 h with 10 μM H_2_O_2_ were consistant with the RNA-Seq data (Table S5).

### 3.5 Transcriptional regulator enrichment analysis

Having confirmed that expression of genes in the NTRK and FOXO clusters was H_2_O_2_-dependent, we next hypothesised that the DEG within each cluster would share common upstream regulatory elements. Identifying these elements could provide new insights into the mechanism(s) by which H_2_O_2_ regulates gene expression. The DEG within the Reactome pathways contributing to the NTRK and FOXO clusters (Table S6) were first analysed with TRAP. TRAP uses the JASPAR database to predict binding motifs based on the 1000 bp upstream flanking region of submitted genes. Analysis with TRAP identified 11 and 15 transcriptional regulators for the FOXO and NTRK clusters respectively (Tables 1 and 2). We next obtained the motifs for these significant hits. Five JASPAR motifs were identified for the FOXO DEG and eight for the NTRK DEG (Tables 1 and 2). JASPAR motifs 21 and 27 (Figure S2) accounted for the majority of identified transcription regulators in both gene sets.

**Table 1:** Potential transcriptional regulators of genes in the FOXO cluster.

| Transcriptional Regulator | TRAP (combined p-value) | JASPAR motif | ChEA3 |  |  |  |  |  |
| --- | --- | --- | --- | --- | --- | --- | --- | --- |
|  |  |  | Encode | Remap | Literature | ARCHS4 | GTE<sub>x</sub> | Enrichr |
| Mycn | 0.0006 | 21 | NA | NS | NS | NS | NS | *** |
| Klf4 | 0.0010 | 27 | NA | NS | ** | NS | NS | *** |
| Egr1 | 0.0013 | 27 | ** | NS | ** | NS | * | *** |
| USF1 | 0.0015 | 21 | ** | NS | NA | NS | NS | *** |
| SP1 | 0.0015 | 27 | * | NS | NA | * | NS | *** |
| Arnt | 0.0017 | 21 | NA | NS | * | * | NS | *** |
| MAX | 0.0017 | 21 | *** | NS | NA | NS | NS | *** |
| znf143 | 0.020 | 63 | NS | NS | NA | * | NS | *** |
| FOXO3 | 0.024 | 10 | NA | NA | ** | ** | ** | *** |
| Myc | 0.028 | 21 | * | NS | ** | NS | * | *** |
| HLF | 0.046 | 5 | NA | NS | NS | NS | NS | NA |

\* $P < 0.05$ , \*\* $P < 0.01$ , \*\*\* $P < 0.001$ . NS, non-significant; NA, not in database. JASPAR matrix IDs and P values for ChEA3 data are in Table S7.

**Table 2:** Potential transcriptional regulators of genes in the NTRK cluster.

| Transcriptional Regulator | TRAP (combined p-value) | JASPAR cluster | ChEA3 |  |  |  |  |  |
| --- | --- | --- | --- | --- | --- | --- | --- | --- |
|  |  |  | Encode | Remap | Literature | ARCHS4 | GTE<sub>x</sub> | Enrichr |
| FOXO3 | 0.00056 | 10 | NA | NA | NS | NS | ** | *** |
| USF1 | 0.0018 | 21 | *** | NS | NA | NS | NS | *** |
| Arnt | 0.0020 | 21 | NA | ** | NS | NS | NS | *** |
| Mycn | 0.0030 | 21 | NA | NS | NS | NS | NS | *** |
| MAX | 0.0037 | 21 | * | * | NA | NS | NS | *** |
| FOXD1 | 0.016 | 10 | NA | NA | NA | NS | NS | NA |
| NFYA | 0.017 | 17 | NS | NS | NA | NS | NS | *** |
| HLF | 0.018 | 5 | NA | NA | NA | NS | NS | *** |
| SP1 | 0.021 | 27 | NS | NS | NA | * | NS | *** |
| RELA | 0.023 | 32 | ** | ** | *** | * | NS | *** |
| Egr1 | 0.024 | 27 | ** | * | *** | *** | *** | *** |
| IRF1 | 0.026 | 42 | NS | NS | NS | NS | NS | *** |
| REL | 0.028 | 32 | NA | NA | NA | NS | NS | *** |
| Myc | 0.030 | 21 | ** | NS | * | NS | NS | *** |
| EBF1 | 0.035 | 7 | NS | ** | NA | NS | NS | *** |

\* $P < 0.05$ , \*\* $P < 0.01$ , \*\*\* $P < 0.001$ . NS, non-significant; NA, not in database. JASPAR matrix IDs and P values for ChEA3 data are in Table S8.

We then compared these results with those obtained from ChEA3, which ranks transcriptional regulators associated with specific gene sets based on ChIPSeq and RNA-Seq data. The six databases within ChEA3 vary in the number of transcriptional regulators represented (118-1628, compared to 841 in JASPAR). Therefore for each transcriptional regulator identified with TRAP, we extracted the *P*_adj_ values across the ChEA3 databases (Tables 1 and 2). This allowed us to determine whether putative H_2_O_2_ responsive transcriptional regulators identified by TRAP could be confirmed with a different analysis method. Reassuringly all significant hits from TRAP were also significant in at least 1, and up to 6, ChEA3 databases. This suggests the identified DNA binding motifs and associated transcriptional regulators are bone fide targets of H_2_O_2_.

A small number of transcriptional regulators identified in the ChEA3 analysis of the NTRK and FOXO gene sets were not present in JASPAR (Tables S9 and 10). In particular, HBP1 and ZBTB10 were significantly enriched from the FOXO gene set, with CSRNP1 and GTF2I significantly enriched from the NTRK gene set, suggesting that these transcriptional regulators may also be responsive to H_2_O_2_.

We also performed TRAP and ChEA3 with the 1050 DEG with fold-change >2, and with a control set of 1050 randomly selected genes. Of the 1050 genes with fold-change >2 the 1000 bp upstream sequence could be mapped for 904 genes for use in TRAP, and 791 had a gene symbol for use in ChEA3. For the 1050 control genes, the 1000 bp upstream sequence could be mapped for all. TRAP analysis of DEG with fold-change >2 identified 32 transcriptional regulators were also significant in one or more of the ChEA3 databases (Table S11). This included 14 that were identified in the analysis of the gene sets associated with the FOXO and/or NTRK Reactome pathway clusters. The 32 transcriptional regulators were represented by 23 JAPSAR clusters, including clusters 21 (16%) and 27 (9%), consistent with the FOXO and NTRK data. Using the control gene set, 38 transcriptional regulators had a significant combined *P* value in the TRAP analysis of the control set of genes. However, none were significant in any of the ChEA3 databases (Table S12). The combined use of the TRAP and ChEA3 data is therefore a strong approach for identifying transcriptional regulators sensitive to H_2_O_2_ or other stimuli.

## 4. Discussion

H_2_O_2_ is produced in a regulated manner in response to a wide range of stimuli. Here we mimiced stimulus-induced H_2_O_2_ production by a plasma membrane NADPH oxidase. By carrying out RNA-Seq followed by pathway and transcription factor motif analyses we have characterised the effect of a physiologically relevant dose of H_2_O_2_ on gene expression, and gained insights into the pathways and transcription factors mediating gene expression changes.

Only a small number of studies have investigated the short term gene expression response to sub-lethal H_2_O_2_ concentrations. An early microarray study identified 828-1097 DEG 5 h after treatment of MCF-7a, MCF-7b or MRC-9 cells with 100 μM H_2_O_2_, with many of these genes related to genotoxic stress (20). Treatment of PIG1 cells with 100 μM H_2_O_2_ resulted in only 661 DEG after 6 h (26), but no analysis was presented on the functional classes of these genes. In HUVEC treated with 10 μM H_2_O_2_, no DEG were identified up to 900 min post-treatment, although 300 μM H_2_O_2_ resulted in maximum DEG (3540) at 4.5 h (29). In contrast we identified 6803 DEG in Jurkat cells treated with a 10 μM bolus dose of H_2_O_2_. Our greater sensitivity is most likely due to the strong statistical power from analysing eight biological replicates compared to three or less in other studies (54). The majority of DEG (85%) have less than 2-fold change in expression and within this set there were similar numbers of up and down regulated genes. For the DEG with a robust >2-fold change in expression, the majority were upregulated. This result suggests that a physiologically relevant concentration of H_2_O_2_ predominantly activates gene expression. There are at least two possible mechanisms by which H_2_O_2_ alters gene expression, and both are likely responsible for our observations. Firstly, H_2_O_2_ is a well described activator of signalling cascades leading to transcription factor activation (2,6,55). Secondly a low dose of H_2_O_2_ decreases DNA methylation 4 h after treatment (56), possibly due to inhibition of DNA methyltransferases during DNA replication (57).

To identify signalling cascades that may be activated by H_2_O_2_ we performed pathway analysis focusing on genes with >2-fold change in expression. Two pathways were convincingly identified, FOXO signalling and NTRK signalling. There are four mammalian FOXO transcription factors, FOXO1, FOXO3, FOXO4, and FOXO6, which are characterised by the presence of a ‘forkhead’ DNA binding domain (58). In response to various stimuli these transcription factors induce the expression of genes involved in a variety of cellular processes (apoptosis, DNA damage repair, cell cycle) (58,59). There is a myriad of evidence that FOXO transcription factors are regulated indirectly by ROS, by modulation of post-translational modifications such as phosphorylation and acetylation (60). For example, Essers et al. (2004) showed that treatment of A14 cells with 20-200 μM H_2_O_2_ led to JNK-dependent phosphorylation of FOXO4, nuclear translocation, and transcriptional activation (61). In the present study, the FOXO transcriptional response was induced using a lower dose of H_2_O_2_. There were two interesting observations: FOXO3 and FOXO4 were upregulated in response to H_2_O_2_ treatment, and TRAP and ChEA3 predicted that FOXO3 was a transcriptional regulator controlling the expression of genes from the FOXO cluster. Previously, it has been shown that FOXO3 is able to upregulate FOXO1 and FOXO4 expression in a positive feedback loop (62). It is possible that FOXO3 is the key transcription factor involved in activating the FOXO transcriptional response to a physiologically relevant dose of H_2_O_2_.

H_2_O_2_ also specifically activated a gene expression signature consistent with NTRK pathway activation. The NTRK genes encode transmembrane tyrosine kinase receptors, TRKA, TRKB, and TRKC (63). Upon binding of their ligands (NGF, BDNF, and NT-3) the TRKA, TRKB, and TRKC receptors dimerise and initiate phosphorylation cascades via PI3K, Ras, and PLCγ (64–66). Our data suggests that one or more of these branches of the NTRK pathway are activated by a low, physiologically relevant dose of H_2_O_2_. Many of the upregulated genes within the NTRK signalling pathway identified in this analysis were MAP kinases, and the transcription factors FOS, FOSB, JUND, and JUNB were also upregulated. It is well-established that heterodimeric AP-1 complexes, made up of FOS and JUN proteins, are both directly and indirectly regulated by concentrations of H_2_O_2_ that would induce oxidative stress (67–69). Both TRAP and ChEA3 predicted that the genes in the NTRK cluster may be activated by RELA/REL, subunits of the NF-κB transcription factor (70). NF-κB was first found to be regulated by H_2_O_2_ in 1991 (71), however, the exact molecular mechanisms are still unclear. A recent study highlighted a link between NTRK signalling and NF-κB activation. Treatment of the KM12SM cell line with NTRK inhibitors, LOXO-101, entrectinib, and dovitinib, led to reduced protein expression of phosphorylated NF-κB (72). Therefore, it is possible that the activation of NF-κB could be leading to regulation of the NTRK signalling pathway.

Taken together, these results suggest that at physiologically relevant levels, H_2_O_2_ acts as a bona fide signalling molecule to stimulate widespread changes in gene expression, with the FOXO and NTRK signalling pathways identified as specific targets. It was perhaps surprising that additional pathways were not identified. However, pathway identification is limited to the extent to which the DEG can be mapped. Indeed only 50% of all DEG, and 37% of DEG with fold-change greater than 2, were available for analysis by Reactome Pathways. Thus, a significant amount of information is lost, and there will be a bias towards well-studied pathways. It is highly likely that H_2_O_2_ activates additional pathways, and as the number of mappable genes increases, reanalysis of the DEG identified in this study will be informative.

