## Supplementary Information for "Insights into H_2_O_2_-induced signaling in Jurkat cells from analysis of gene expression"

### Supplementary Data

**Figure S1:** Principal component analysis (PCA) of the RNA-Seq data

**Figure S2**: Root motif for Jasper clusters 21 and 27

**Table S1:** RT-qPCR Primers

**Table S2:** Significantly differentially expressed genes between WT and WT treated cells

**Table S3:** List of 21 enriched Reactome pathways for mapped DEG with >2-fold change in expression

**Table S4:** List of 351 enriched Reactome pathways for all mapped DEG

**Table S5:** Comparison of fold change (FC) values from qPCR and RNA-Seq 4 h after treatment with 10 µM $H_2O_2$

**Table S6:** FOXO and NTRK cluster DEG used for motif analysis

**Table S7:** JASPAR Matrix ID and ChEA3 data p-values for transcriptional regulators identified by TRAP (combined $p<0.05$) for genes in the FOXO cluster

**Table S8:** JASPAR Matrix ID and ChEA3 data p-values for transcriptional regulators identified by TRAP (combined $p<0.05$) for genes in NTRK cluster

**Table S9:** ChEA3 Mean Rank output for genes from the FOXO cluster

**Table S10:** ChEA3 Mean Rank output for genes from the NTRK cluster

**Table S11:** TRAP, JASPAR clusters, and ChEA3 output for 1050 genes with L2FC > 1

**Table S12:** TRAP, JASPAR clusters, and ChEA3 output for 1050 control genes

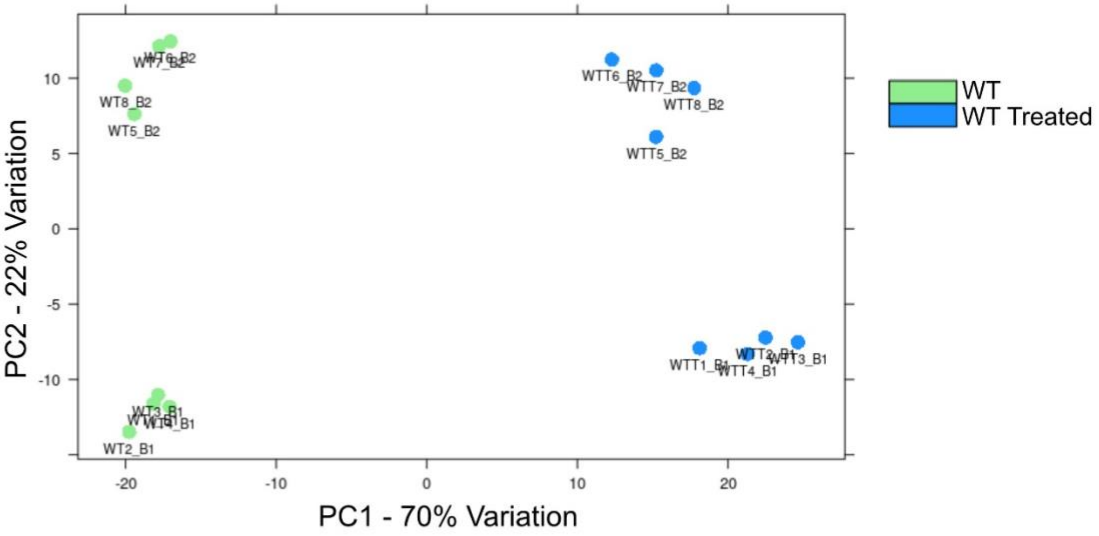


**Figure S1. Principal component analysis (PCA) of the RNA-Seq data based on 16 independent samples.** There was distinct clustering of treated samples (WTT - blue dots) and untreated samples (WT - green dots) within the first principal component, explaining 70% of the variation. The second principal component represented the batch effect from independent library preparations and sequencing in two batches (B1 vs B2).

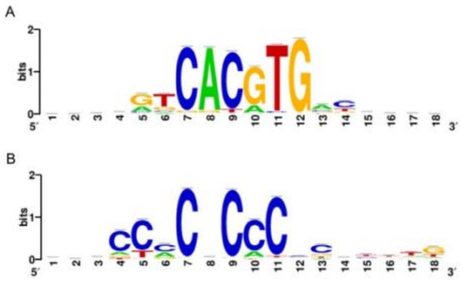


**Figure 6**. **Root motif for Jasper cluster 21 (top) and 27 (bottom).**
